# Emerin expression stratification across breast cancer subtypes

**DOI:** 10.1101/2025.07.03.663032

**Authors:** Thaysa Ghiarone, Emily Hansen, James M. Holaska

## Abstract

Nuclear dysmorphism is a critical indicator of tumor aggressiveness, influencing cancer cell invasion and metastasis. Emerin, an integral nuclear envelope protein involved in nuclear architecture, is important for maintaining nuclear integrity. Our previous work demonstrated an inverse correlation between nuclear envelope-localized emerin expression and breast cancer aggressiveness. However, it failed to have the power to assess whether emerin loss correlates with cancer stage, grade, proliferation, or molecular phenotype. Here we analyzed emerin expression at the nuclear envelope across 243 breast cancer patient samples encompassing various tumor grades, stages, and molecular phenotypes. We found significantly reduced emerin expression in invasive ductal carcinoma (IDC), invasive lobular carcinoma (ILC), and ductal carcinoma in situ (DCIS), compared to normal breast tissue. Notably, emerin loss correlated with advanced tumor stage, higher Ki-67 proliferation rates, elevated human epidermal growth factor receptor 2 (HER2) levels, and decreased estrogen receptor (ER) and progesterone receptor (PR) expression—markers associated with more aggressive breast cancers. Emerin expression was consistently reduced in triple-negative breast cancer (TNBC) and other receptor-negative subtypes, underscoring its potential role in tumor dedifferentiation and progression. These findings highlight emerin as a promising prognostic biomarker and therapeutic target for aggressive breast cancer subtypes.

## Introduction

Nuclear morphology is a well-established diagnostic tool for grading cancer,^1^ with changes in nuclear shape and compliance serving as key indicators of tumor aggressiveness.^2–4^ One recognized ‘hallmark of cancer’ is nuclear softening,^5, 6^ which facilitates cancer cell movement through tissues and endothelial slits, promoting invasion and metastasis.^7^ This nuclear softening is influenced by tumor microenvironment (TME) stiffness, which is driven primarily by increased extracellular matrix (ECM) deposition by the tumor.^8^ Notably, stiffening of these invasive, malleable breast cancer cell nuclei can reduce their invasion and metastatic potential,^9^ highlighting nuclear stiffness as a crucial factor in cancer progression.

Nuclear stiffness is regulated by a network of nucleostructural proteins that serve both as structural scaffolds and signaling mediators. Emerin, an inner nuclear membrane protein that interacts with lamins and actin, plays a key role in nuclear architecture.^10–12^ Mutations in emerin, particularly within its nucleoskeletal binding domain, have been identified in various cancers.^11^ Our previous studies demonstrated that triple-negative breast cancer (TNBC) cell lines exhibit significantly reduced emerin expression compared to non-cancerous controls, correlating with smaller nuclear size, nuclear dysmorphism, increased migration, and enhanced invasion.^12, 13^ Reintroducing GFP-tagged wild-type emerin rescued these defects, whereas mutant emerin forms incapable of binding nuclear actin and lamins failed to restore nuclear structure and function.^12^ In vivo, wild-type GFP-emerin expression in MDA-MB-231 cells reduced primary tumor size and lung metastasis, whereas emerin knockdown in MCF7 cells promoted migration and nuclear deformity,^13^ further supporting a role for emerin loss in cancer progression. Importantly, emerin expression at the nuclear envelope was inversely correlated with tumor aggressiveness and metastasis in patient samples.^13^

Emerin dysregulation extends beyond breast cancer, with studies implicating its loss in prostate, hepatocellular, lung, and ovarian cancers.^14–16^ In prostate cancer, emerin inhibits metastasis, and its loss is linked to increased nuclear deformity and migration.^14^ Recent studies also show that emerin loss is associated with increases in stem-like activity of prostate cancer cells.^17^ Similarly, hepatocellular carcinoma (HCC) cells with reduced emerin expression exhibit enhanced migration and invasion.^15, 16^ In ovarian cancer, decreased emerin levels were observed in 38% of cases and were associated with nuclear envelope defects, impaired nuclear reorganization, and epithelial-to-mesenchymal transition (EMT).^18–20^ These findings suggest that emerin reduction may drive invasive phenotypes across multiple cancer types.

Our previous work demonstrated an inverse correlation between nuclear envelope-localized emerin expression and breast cancer aggressiveness,^13^ though additional patient samples were needed to assess whether emerin loss correlates with cancer stage, grade, proliferation, or molecular phenotype. In this study, we analyzed over 200 patient samples across all breast cancer grades and molecular subtypes to further investigate these associations.

We show that reduced emerin expression correlates with increased Ki-67 positivity, elevated HER2 expression, and decreased ER and PR expression—features associated with more aggressive tumors. Additionally, emerin loss is linked to cancer stages involving local lymph node metastasis and beyond. Interestingly, emerin reduction was consistent across all tumor grades, suggesting that loss of emerin may contribute to cellular dedifferentiation. These findings support further investigation into emerin as a potential biomarker for identifying tumors prone to invasion and metastasis, which may benefit from more aggressive treatment strategies.

## Results

We analyzed emerin expression on 86 additional patient samples by immunohistochemistry with emerin antibodies (10351-1-AP, Proteintech) on a tissue microarray (BR804b and BR480b, TissueArray, LLC) to increase the power of our studies. This array contained breast cancer tumor samples from a range of types, stages, and grades. Each tumor was assessed for emerin expression at the nuclear envelope (NE grade) based on the amount of emerin staining at the nuclear periphery; this accounts for both protein expression and normal localization.^13^ The grader was blinded to sample identifiers. After excluding tissue samples that were unable to be analyzed (i.e., damaged or folded tissues), we were able to analyze 84 samples. Similar to our previous findings, emerin protein is abundant at the nuclear envelope in normal breast tissue (2.0 ± 0.22) and in breast hyperplasia (2.1 ± 0.27), while emerin expression at the nuclear envelope was significantly lower in ductal carcinoma in-situ (DCIS; 1.173 ± 0.6), invasive ductal carcinoma (IDC; 0.895 ± 0.66), and invasive lobular carcinoma (ILC; 0.619 ± 0.49) in these new 86 patient samples (**Figure 1A,B**). To confirm that these results were emerin-specific, we stained this tissue microarray with just secondary antibody to account for background staining (**Figure 1A, bottom right panel)**.

**Figure 1:**
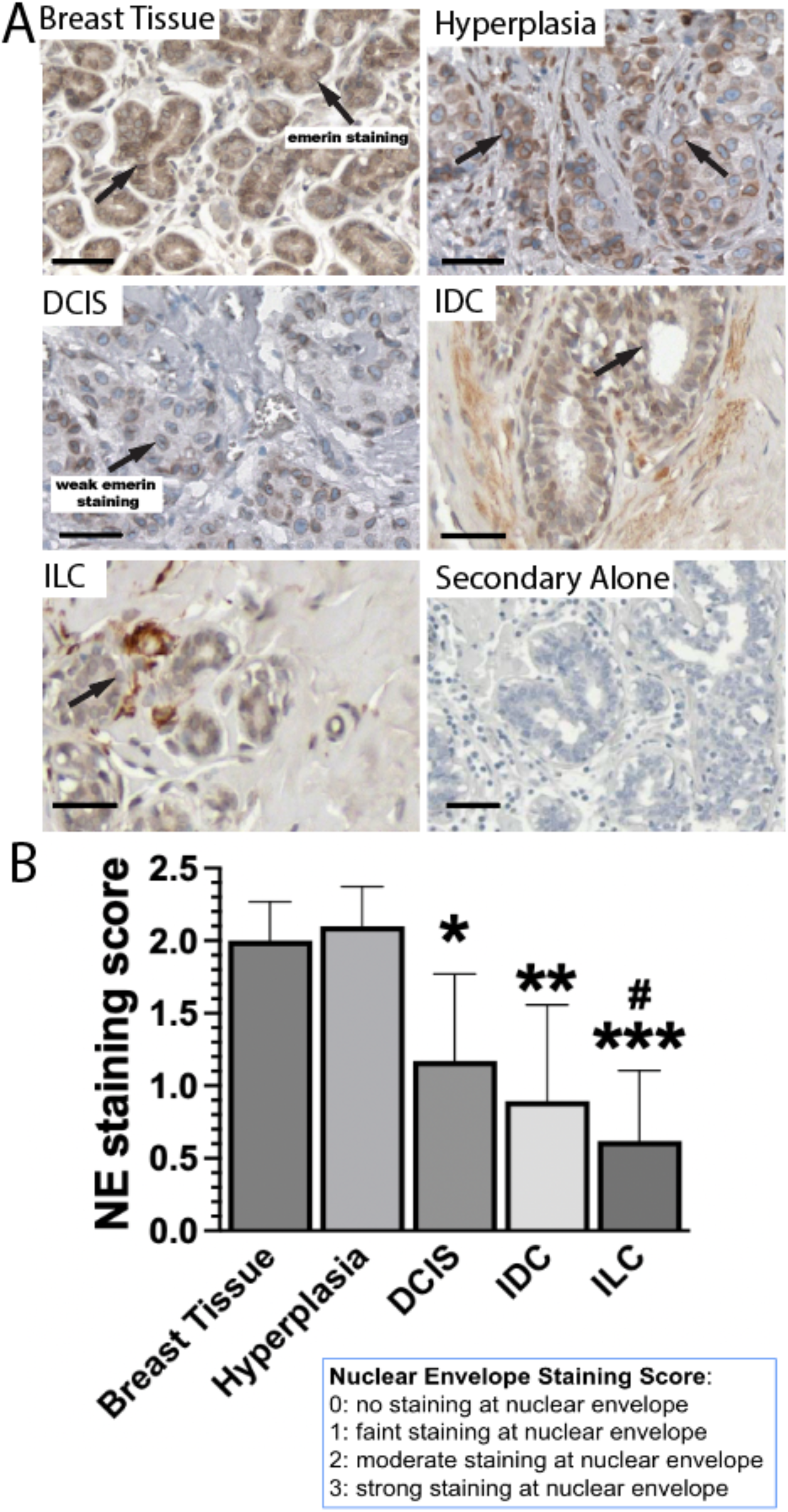
Reduced emerin expression at the nuclear periphery correlates with breast cancer invasiveness in patients. A) Representative tissue microarray staining of emerin in tissues of 84 patients using emerin polyclonal antibodies (Proteintech, cat# 10351-1-AP) or secondary alone (Vector Lab, cat#: MP-7451). Nuclei are blue and emerin is brown. As severity of cases increases, there is a visible reduction in emerin expression at the nuclear envelope and more deformed nuclei are present. B) Quantification of emerin staining on IHC-stained patient samples using a scale of 0-3, with 0 having no staining at the nuclear periphery and 3 having complete, dark rim staining. DCIS= ductal carcinoma in-situ, IDC= invasive ductal carcinoma, ILC= invasive lobular carcinoma. N=84 total samples. ***P≤0.0003 comparing breast tissue and hyperplasia to ILC, **P≤0.0016 comparing breast tissue and hyperplasia to IDC, *P≤0.0455 comparing breast tissue and hyperplasia to DCIS, and ^#^P=0.0201 comparing DCIS to ILC, one-way ANOVA followed by Tukey’s Test. Error bars represent standard deviation. Scale bar = 50 μm.

These samples were combined with samples from BR2082c (TissueArray) for a total of 243 patient samples that were of sufficient quality to be analyzed and had relevant grading, staging, and molecular phenotyping. Grades 1 (1.168 ± 0.96), 2 (1.213 ± 0.79), and 3 (1.18 ± 0.41) all had significantly decreased emerin expression at the nuclear envelope when compared to non-cancerous samples (2.69 ± 0.446) and tumor samples with no grade (2.31 ± 0.67), with p<0.0001 for each (**Figure 2A,B**). It was found that stage IIIA tumors had the lowest emerin expression (0.544 ± 0.67), followed by stages IIIB (0.836 ± 0.44) and IIB (0.955 ± 0.65), with p=0.0004, p=0.0422, and p=0.008, respectively, when compared to stage 0 (1.75 ± 0.84; **Figure 2C,D**). Stages IA (1.73 ± 1.02), and IIA (1.41 ± 0.89) were not significantly different. Non-cancerous samples were NE graded 2.69 ± 0.45 and significantly higher than all other samples (p<0.024).

**Figure 2:**
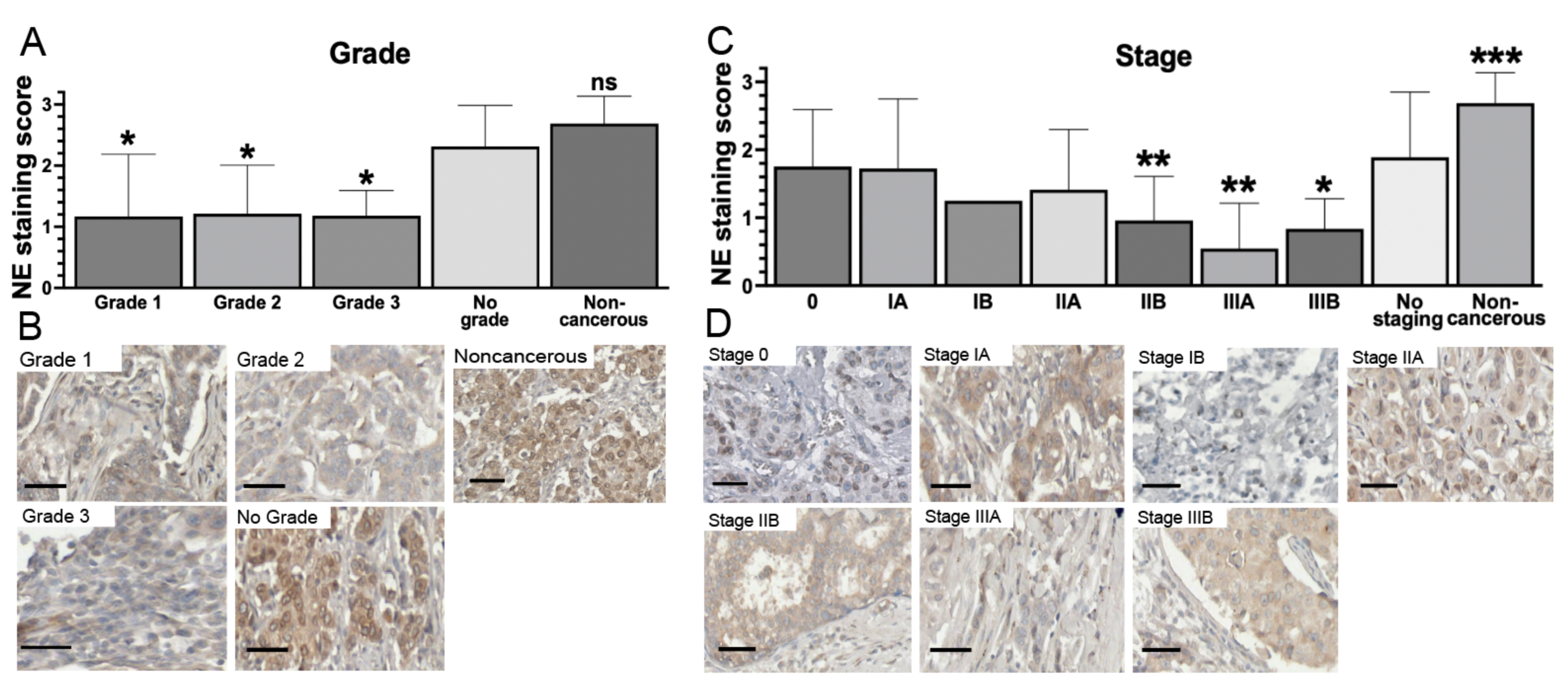
Increased grading and staging of cancer correlates with decreased emerin expression at the nuclear periphery. A, C) Quantification of emerin nuclear envelope staining on patient samples using a 0-3 scale, with 0 having no staining at the nuclear periphery and 3 having complete, dark rim staining. B, D) Representative tissue microarray staining of emerin in 243 patients using emerin polyclonal antibody (Proteintech, cat# 10351-1-AP). Nuclei are blue and emerin is brown. N=243 samples. A) *P<0.0001 compared to noncancerous, NS= not significant between no grade and noncancerous. C) ***P<0.0001, **P≤0.008, *P=0.0422 with one-way ANOVA followed by Dunnett’s test for comparing all samples to stage 0. Error bars represent standard deviation.

To examine the relationship between emerin expression and molecular phenotype, we compared emerin nuclear envelope levels to ER, PR, and HER2 expression, as well as Ki-67 positivity to denote proliferation. Tumors lacking ER expression had the lowest emerin expression (neg, 1.139 ± 0.73, p<0.0001), while ER levels graded as + (1.973 ± 0.106, p=0.0223), ++ (1.66 ± 1.04, p=0.0001), and +++ (1.47 ± 0.92, p<0.0001) had more emerin expression at the nuclear envelope (**Figure 3A,B**). Note that p-values are relative to non-cancerous samples. There was a significant difference between ER-negative samples and ER + samples, with p=0.0019. Tumors lacking PR expression had the lowest emerin expression (neg, 1.159 ± 0.78), while PR levels graded as + (1.601 ± 1.11), ++ (1.76 ± 1.11), and +++ (1.553 ± 0.89) had more emerin expression at the nuclear envelope (**Figure 3C,D**). P≤0.0004 for all samples relative to non-cancerous samples. There was a significant difference between PR-negative samples and PR ++ samples, with p=0.0145. Tumors lacking HER2 expression (0, 1.683 ± 1.02; **Figure 3E,F**) did not have statistically different emerin expression at the nuclear envelope than those graded as HER2 1+ (1.038 ± 0.92) or 2+ (1.301 ± 1.016). However, compared to HER2 0, HER2 3+ (1.031 ± 0.607) was significantly lower (p<0.0001). All of them were all significantly (p≤0.0034 for all samples) lower than non-cancerous control (2.69 ± 0.446). Interestingly, emerin NE expression in TNBC samples were not statistically different than any of the double-negative molecular phenotypes (**Figure 3G,H**), suggesting that decreased expression of any combination of two of ER, PR, and HER2 is sufficient to reduce emerin expression.

**Figure 3:**
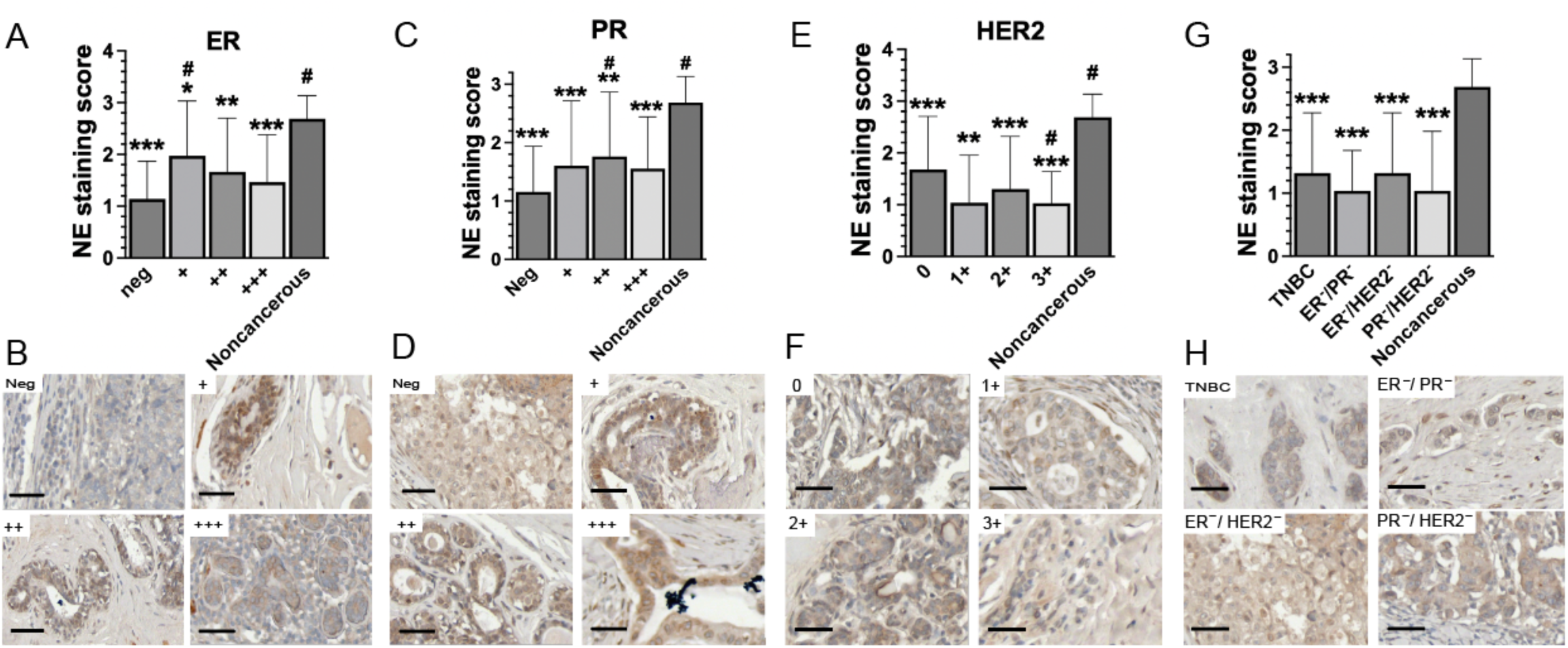
Lack of ER expression, PR expression, or a combination of decreased ER, PR, and HER2 expression is correlated with decreased emerin expression at the nuclear periphery. Quantification of emerin expression at the nuclear periphery relative to levels of A) ER, C) PR, and E) HER2 expression in breast cancer and B, D, F) representative tissue microarray staining, respectively, using an emerin polyclonal antibody (Proteintech, cat# 10351-1-AP). G) quantification of emerin expression at the nuclear periphery comparing noncancerous tissue to cancerous tissue lacking a combination of PR/HER2, ER/HER2, ER/PR, or all three receptors (TNBC) and H) representative images, respectively. Samples were graded using a scale of 0-3, with 0 having no staining at the nuclear periphery and 3 having complete, dark rim staining. N=243 samples. A,C,E,G) ***P<0.0001, **P≤0.004, *P=0.0223 compared to noncancerous within graphs. A) ^#^P≤0.023 compared to neg, C) ^#^P≤0.0145 compared to negative, E) ^#^P<0.0001 compared to HER2 0, one-way ANOVA followed by Tukey’s test.

Emerin expression at the nuclear envelope inversely correlated with Ki-67 expression and thus proliferation. Ki-67 negative tumors had similar emerin expression (2.298 ± 0.95; **Figure 4A,B**) as compared to the non-cancerous samples (2.69 ± 0.446). Tumors that were 1-9% Ki-67-positive had a NE score of 1.452 ± 0.91 (p<0.0001). Tumors that were 10-19% Ki-67-positive had a NE score of 1.201 ± 0.81 (p<0.0001). Tumors that were 20-49% Ki-67-positive had a NE score of 0.9111 ± 0.58 (p<0.0001) and tumors that were >50% Ki-67-positive had a NE score of 0.7688 ± 0.42 (p<0.0001). Note all of these p-values are relative to non-cancerous samples. When comparing to 1-9% Ki-67- positive samples to all categories of Ki-67 expression, 1-9% Ki-67-positive samples were significantly different from all other categories (p<0.0001).

**Figure 4:**
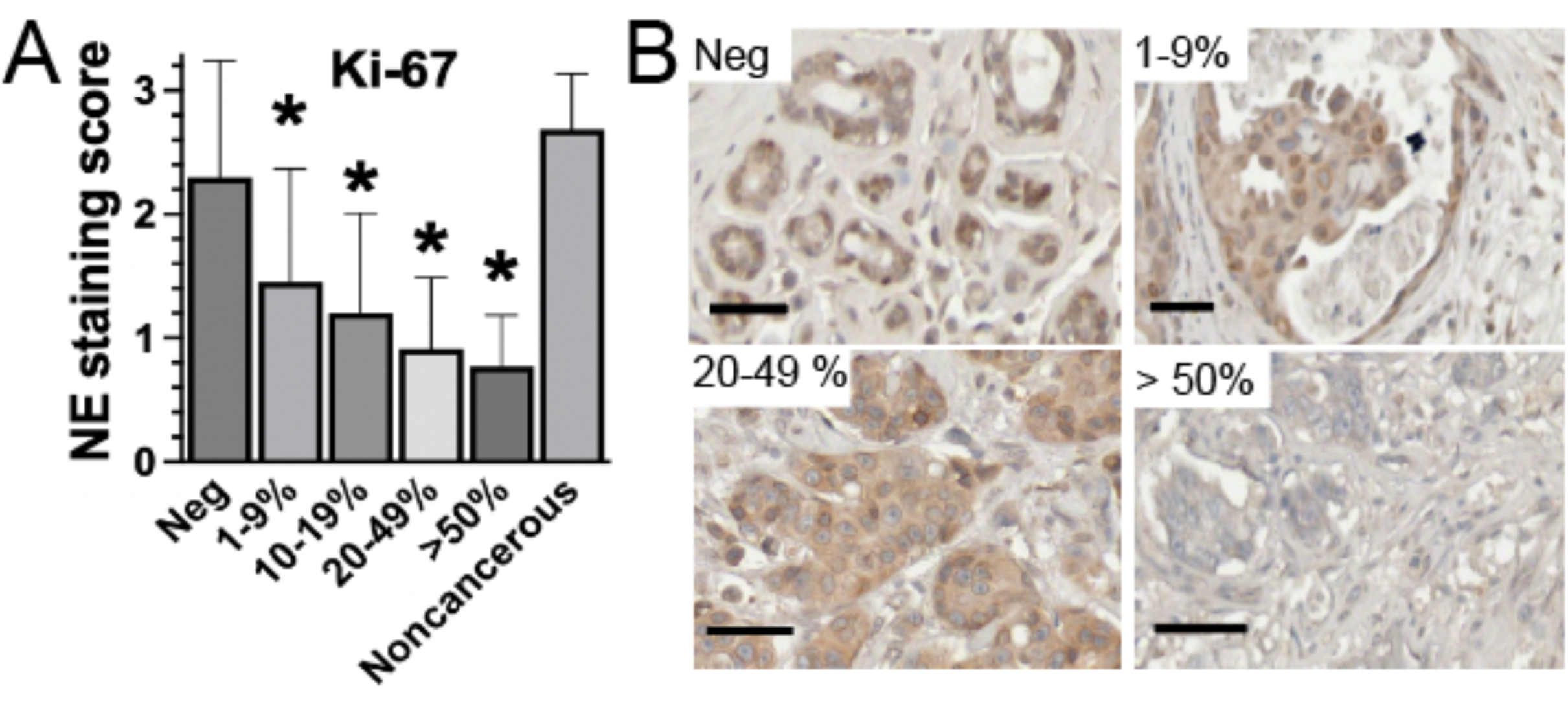
Emerin expression at the nuclear envelope inversely correlates with Ki-67 expression. A) Quantification of emerin expression at the nuclear periphery relative to the percentage of Ki-67 staining and B) representative images of cancerous Ki-67 negative, 1-9%, 20-49%, and >50% Ki-67-stained tissue. N=243 samples. *P<0.0001 compared to Ki-67 negative and noncancerous tissue, one-way ANOVA followed by Tukey’s test. Error bars represent standard deviation.

## Discussion

For decades the presence of abnormal nuclear structure was used to distinguish tumor cells from normal cells, and to grade tumors.^11, 21, 22^ Our findings reinforce the role of nuclear morphology as a critical diagnostic and prognostic tool in breast cancer. Previous studies have established that nuclear deformability and softening contribute to increased tumor invasiveness and metastasis, primarily by facilitating cell migration through the extracellular matrix and vasculature.^11–13, 23, 24^ Our results demonstrate that emerin expression at the nuclear envelope is significantly reduced in breast cancer patient samples, particularly in invasive ductal and lobular carcinomas, further implicating nuclear envelope integrity in cancer progression.

Thus, we conclude that loss of emerin is crucial for transforming benign tumor cells to a more invasive phenotype. Consistent with our results, reduction in emerin also correlated with nuclear softening in melanoma cells,^25^ something that is well-established to correlate with rates of invasiveness and metastasis.^26, 27^

### Emerin Expression and Breast Cancer Stratification

We found that emerin expression at the nuclear envelope is inversely associated with breast cancer aggressiveness. Specifically, emerin expression decreases in a stepwise manner from normal breast tissue to hyperplasia, DCIS, and then invasive breast cancers. This progressive loss of emerin is consistent with its hypothesized role in maintaining nuclear stability and suppressing invasive properties in epithelial cells.

Furthermore, our data reveal that lower emerin expression is associated with higher tumor grade and more advanced cancer stage, with stage IIIA tumors exhibiting the lowest emerin levels. These findings suggest that emerin loss occurs early in tumor progression and continues to decline as cancer advances, which may contribute to increased nuclear deformability and enhanced metastatic potential.

These results support a model by which emerin downregulation occurs in a cell population within a growing tumor. These cells would then be selected during tumor evolution because of their increased proliferation and increased nuclear compliance, which allows them to be more invasive. This increased invasiveness enables the cells to invade the extracellular matrix and squeeze through the vascular endothelium to promote increased cancer cell survival and metastasis. Supporting our results, recent studies in prostate^14^ and ovarian cancer,^19^ found that patients have decreased emerin and that this decreased emerin contributes to higher nuclear deformity and invasion.^14^

### Emerin and Breast Cancer Molecular Phenotyping

Stratification of tumors based on ER, PR, HER2, and Ki-67 expression is a cornerstone of breast cancer diagnosis and treatment planning.^28–31^ Our results show that emerin expression at the nuclear envelope is significantly reduced in ER-negative and PR-negative tumors compared to ER-positive and PR-positive tumors. Given that ER and PR positivity generally correspond to better prognoses and more differentiated tumors,^30^ our findings support the hypothesis that emerin plays a role in maintaining cellular differentiation and nuclear integrity. The inverse correlation between emerin and Ki-67, a marker of proliferation,^32^ further reinforces this concept, as lower emerin expression is linked to increased tumor cell proliferation, a key characteristic of aggressive cancers.

Although HER2 expression alone did not show a strong correlation with emerin expression, the triple-negative breast cancer (TNBC) subtype, which lacks ER, PR, and HER2 expression, exhibited similarly reduced emerin levels as double-negative phenotypes. This suggests that the loss of any combination of ER, PR, and HER2 is associated with decreased emerin, reinforcing the idea that emerin loss may be a marker of dedifferentiation and aggressive tumor behavior.

### Implications for Emerin as a Biomarker

The strong correlation between reduced emerin expression and increased tumor aggressiveness, proliferation, and loss of differentiation markers suggests that emerin could serve as a valuable prognostic biomarker in breast cancer. The ability of emerin expression to distinguish between less aggressive and highly invasive tumor types indicates that it may help identify patients who are at higher risk of progression and metastasis. Furthermore, the relationship between emerin and Ki-67 expression suggests that emerin loss may mark tumors with higher proliferative capacity, which are more likely to respond to aggressive treatment regimens.

Given the emerging evidence linking emerin to nuclear mechanotransduction, chromatin organization, and gene regulation,^33^ further studies should explore the molecular mechanisms by which emerin modulates breast cancer progression. Investigating whether restoring emerin expression can suppress invasiveness and metastasis may open new avenues for targeted therapies aimed at reinforcing nuclear stability in aggressive breast cancers.

## Conclusion

This study provides compelling evidence that emerin expression at the nuclear envelope is an important factor in breast cancer progression. Its inverse correlation with tumor grade, stage, ER/PR expression, and proliferation markers highlights its potential as both a diagnostic and prognostic tool. Future research should aim to elucidate the mechanistic role of emerin in nuclear architecture and cancer cell invasion to determine its viability as a therapeutic target.

By further investigating emerin’s role in breast cancer progression, targetable treatments for even the most invasive breast cancer types, such as triple-negative breast cancer, may be revealed. The observed negative relationship between emerin expression and advanced breast cancer suggests that restoring emerin function could serve as a treatment, particularly in advanced stages of cancer that lack effective treatment options.

## Materials and methods

### Immunohistochemistry and Tissue Microarray Analysis

Tissue microarrays (TissueArray, LLC, cat# BR2082c for emerin-stained samples and cat# BR087e094 for secondary-only tissue) were deparaffinized and rehydrated in Coplin jars (4 minutes in xylene three times, 3 minutes in 100% EtOH two times, 3 minutes in 95% EtOH two times, 3 minutes in 80% EtOH, 3 minutes in 70% EtOH, and 5 minutes in distilled water) prior to being placed in citrate buffer (0.05% Tween 20, 10mM citric acid at pH 6.0) and steamed at 95°C for 45 minutes. Slides were cooled at RT and washed with PBS. Endogenous peroxidase was removed by washing with 0.3% hydrogen peroxide for 20 minutes at room temperature in coplin jars. After washing again with PBS, slides were blocked in 1% Bovine Serum Albumin (VWR, cat#: 97061-420) in PBS for 15 minutes. Slides were then incubated with anti-emerin antibody (Proteintech, cat#: 10351-1-AP, 1:500 dilution) for two hours at 37°C in a humidified chamber. Slides were washed again with PBS and blocked with 2.5% Normal Goat Serum for 20 minutes at room temperature (ImmPRESS Reagent, Vector Lab, cat#: MP-7451) and then incubated with the anti-rabbit ImmPRESS IgG peroxidase reagent (Vector Lab, cat#: MP-7451) or the anti-mouse ImmPRESS IgG peroxidase reagent (Vector Lab, cat# MP-7452) per manufacturer instructions. Slides were washed again with PBS and incubated with the ImmPACT DAB peroxidase substrate (Vector Lab, cat#: SK4105) for 90 seconds at room temperature while checking color development under a microscope before rinsing in tap water. Slides were counterstained with Vector Hematoxylin (Gill’s Formula, Vector Lab, cat#: H3401) for three minutes. After rinsing again in tap water, slides were incubated in 0.1% sodium bicarbonate for one minute, rinsed with tap followed by distilled water, and then dehydrated and mounted with Prolong Diamond Antifade Mountant (Invitrogen, cat#: P36970). Tissue images were taken using the Evos FL Auto microscope and the Precipoint slide scanning microscope. Blinded grading of tissues was done using a grading system in which a 0 corresponded to no emerin staining at the nuclear periphery and a 3 corresponded to complete, dark staining of emerin at the nuclear periphery. All grading was done in one sitting to avoid multi-day bias. Invasive, DCIS, and metastatic tissue measurements were determined significant against noncancerous tissue using one-way ANOVA with Tukey or Dunnett’s comparison. Descriptive statistics are presented as Mean ± SD.

## Acknowledgements

We thank the Department of Biomedical Sciences at Cooper Medical School of Rowan University for providing funding for this work and many fruitful discussions. We thank Dr. Isabelle Mercier (St. Joseph’s University) for many fruitful discussions regarding these studies. We thank the members of Holaska’s lab for the numerous discussions pertaining to this manuscript and the Boehning lab (Cooper Medical School at Rowan University) for technical help as needed during experiments.

## Author Contributions

J.M.H., T.G.S., and E.H. conceived the project. T.G.S. and E.H. performed the IHC experiments and T.S.G., E.H., and J.M.H. analyzed the data in the IHC experiments. All authors contributed input in figure preparation and in writing of the manuscript.

## Data Availability

The data supporting the findings of this study are available by request from the corresponding author, J.M.H.

